# Inverted encoding models estimate sensible channel responses for sensible models

**DOI:** 10.1101/642710

**Authors:** Thomas C. Sprague, Geoffrey M. Boynton, John T. Serences

## Abstract

In a commentary published in *eNeuro*, Gardner & Liu (2019) discuss the role of model specification in interpreting the output of complex models of neural data. As a case study, they suggest that one variant of such analyses, the inverted encoding model (IEM) analysis framework, should not be used to assay properties of “stimulus representations” because the ability to apply linear transformations at various stages of the analysis procedure renders results ‘arbitrary’. As we discuss, the *specification* of all models is arbitrary to the extent that an experimenter makes choices based on current knowledge of the model system. However, the *results* derived from any given model, such as the reconstructed channel response profiles obtained from an IEM analysis, are uniquely defined and are arbitrary only in the sense that changes in the model can predictably change results. Moreover, with knowledge of the model used for IEM analyses, the results remain informative as comparisons between reconstructed channel response profiles across task conditions using a fixed encoding model – the most common use of the IEM technique – can generally capture changes in population-level representation magnitude across linear transformations. Thus, changes in the magnitude of the response profiles across conditions are preserved, even across unprincipled linear transforms. IEM-based channel response profiles should therefore not be considered arbitrary when the model is clearly specified and guided by our best understanding of neural population representations in the brain regions being analyzed. Intuitions derived from this case study are important to consider when interpreting results from all model-based analyses, which are similarly contingent upon the specification of the models used.

## Introduction

In any model-based analysis framework, the modeling choices made by the researchers critically influence results of the modeling procedures. That is - it is impossible to interpret the results without knowledge of the details of the model used to generate those results. Moreover, altering the properties of the model should naturally change aspects of the results, sometimes in a predictable and straightforward way. This is true for all model-based analysis frameworks, including the popular single-voxel population receptive field (vRF) modeling approach (Dumoulin and Wandell, 2008; Wandell and Winawer, 2015; Vo et al., 2017b), fitting extremely high-dimensional voxel-wise encoding models to densely-sampled datasets (Kay et al., 2008; Naselaris et al., 2009; Nishimoto et al., 2011; Huth et al., 2012; Çukur et al., 2013b; Huth et al., 2016; Lescroart and Gallant, 2019), the inverted encoding model (IEM) technique (Brouwer and Heeger, 2009, 2011, 2013; Scolari et al., 2012; Foster et al., 2015), and even fitting standard GLMs to fMRI data (Friston et al., 1994).

Typically, the IEM technique involves experimenters estimating a simplified model built of stimulus-selective feature channels (for orientation; color; motion direction; spatial position; polar angle), each tuned to specific feature values and tiling the full stimulus space (Freeman and Adelson, 1991). For example, one could build a model with 8 channels tuned to different stimulus orientations (Brouwer and Heeger, 2011; Ho et al., 2012; Scolari et al., 2012). The properties of these channels are typically inspired by our understanding of the visual system - there are populations of cells tuned to particular orientations; colors; motion directions; positions – and at least in early sensory areas, much is known about the characteristic shape of single-unit tuning functions. Then, based on these modeled channels, linear regression is used to estimate how such a model accounts for changes in activation in a given voxel across different stimulus conditions (fitting the ‘forward’ model). The best-fit model can then be inverted to infer the activation of each modeled channel - that is, the reconstructed channel response profile - given the previously-estimated model and new measured activation patterns across many voxels. The result, when channels are modeled as selective for a single stimulus value, is typically a channel response profile with a peaked response at the feature value(s) present in the stimulus. Importantly, the inversion step effectively summarizes the results by transforming modulations across all measured units (e.g. voxels/EEG electrodes) back into the model space. This is not the only means of aggregating information, but one of a family of approaches to interpret modulations at the level of large-scale activation patterns.

While channel response profiles^1^ often look qualitatively similar to neural tuning functions for single units, a point brought up by Gardner & Liu (2019), it is critical to understand that reconstructed channel response profiles cannot be used to draw conclusive inferences about any specific attributes of single-unit response properties (e.g. single unit gain or bandwidth changes, for more on this, see Sprague et al., 2018). Moreover, recovery of peaked channel response profiles does not demonstrate the accuracy (or inaccuracy) of the channel shapes in the encoding model used (Sprague et al., 2018a; Gardner and Liu, 2019).

In their commentary, Gardner & Liu (2019) argue that the channel response profiles resulting from the IEM technique are “arbitrary” because invertible linear transforms of the basis set will fit the data equally well. Hence, changing the model can predictably change the model fit, which in turn changes the shape of the reconstructed channel response profiles. This ability to apply invertible linear transforms means that any reported channel response profile’s shape is one from an infinite family of shapes (spanned by all invertible linear transforms that could be applied to the analysis). In their words, “the channel response function is only determined up to an invertible linear transform. Thus, these channel response functions are arbitrary, one of an infinite family and therefore not a unique description of population representation.” (Gardner and Liu, 2019; abstract). Thus, if a researcher used an unprincipled set of assumptions about the shape of the modeled channels – that is, ignoring known properties of visual selectivity – then these assumptions can be recapitulated in the reconstructed channel response profiles. For example, Gardner and Liu (2019) showed that if orientation channels are presumed to be bimodal then the resulting reconstructed channel response profiles can also have a bimodal shape. Below we argue that all models are arbitrary, even those informed by biology, but the results of a model are not arbitrary once the model has been specified – this is true for both forward encoding models and inverted encoding models. Next, we show that even if poorly motivated models are used (or poorly motivated linear transforms are applied), differences between conditions are generally preserved. Finally, we discuss important considerations when interpreting IEM-based analyses and what we see as the place for this modelling approach in the context of other useful analysis methods.

## IEM-based channel response profiles are uniquely determined given a fixed model

It is an unfortunate mischaracterization to imply that IEM-based results are “arbitrary” without specifying that they are uniquely determined and interpretable with knowledge of the modeled basis. Although one can generate many descriptions of a population representation, the result is not “arbitrary” if the channel response profile is interpreted in the context of the model used by the researchers. As a simple example, one invertible linear transform that could be applied to a basis and the resulting channel response profiles would ‘shift’ the columns of the predicted channel response matrix by one. This would result in each channel being mislabeled, but all other features of the analysis would proceed intact. With knowledge of this mislabeling (that is, knowledge of the original basis and the invertible linear transform), it is possible to ‘undo’ the transform and to achieve the intended understanding. Likewise, if the experimenter reports their basis (as all IEM reports do, so far as we know), and the reconstructed channel responses or derived measures are computed in the context of that basis, then there are no concerns as to the arbitrariness of the channel response profile’s shape. Thus, even though it is true that if you change certain aspects of your model, you can predictably change aspects of your results, it does not follow that the reconstructed channel response profile is arbitrary – it is simply influenced by the choice of the model, just like the result of any model is influenced by its specification. So, rather than interpreting IEM results as *<u>the</u>* population representation, it is more appropriate to consider them one possible depiction of a population representation, as uniquely derived from a particular model.

## Results from all models depend on properties of the model

Importantly, the points the authors raise about applying invertible linear transforms (that is, changing the coordinate system of a linear model) apply to nearly all model-based analyses, even those that only compute a *forward* encoding model to predict responses of measured neural signals based on stimulus properties, without any attempt at “inversion” back into a stimulus-referred space. We consider two trivial examples: spatial receptive fields measured via single-unit electrophysiology, and a general linear model (GLM) fit to a 2-condition fMRI experiment. When estimating the spatial RF of a neural measurement (either voxel or neuron), it is necessary to relate the observed neural response to changes in the stimulus. Under certain noise assumptions, one could even weight the stimulus aperture (in screen coordinates) by the observed neural signal. But even this procedure involves an implicit set of model assumptions, namely, that the ‘basis’ for the stimulus model is in visual field coordinates (1 number for each location in the visual field). Thus, the same logic of coordinate transforms applies here: one could apply any number of invertible linear transforms to the image basis and to the estimated RF profile, and the resulting model would account for the same amount of variance because it is a linear transform of the original model. For instance, a 2D Fourier transform could be used to losslessly transform between a spatial basis and a Fourier basis. Does this mean we should consider RF (or feature tuning) models estimated in a similar way as arbitrary? Of course not. The existence of a potential coordinate transform does not render the original model invalid, it just means that one must know the model in order to interpret the results.

A similar logic applies to a simple 2-condition fMRI experiment using univariate statistical approaches (i.e. voxel-wise analysis with a GLM; Friston et al., 1994). Consider the case where a participant is sometimes pressing a left button and sometimes a right button. The experimenter can build a GLM with predictors for BOLD activation associated with pressing the left and right button, appropriately convolved with a model hemodynamic response function. In turn, the experimenter could apply the invertible linear transform P = [0 1; 1 0] to the model basis (and thus, the resulting GLM regressors), which would result in flipped estimated beta weights: the beta weight originally corresponding to right now corresponds to left, and vice versa. But, of course, you know the ‘original’ layout of the regressors, so you could just update your labels of the weights accordingly. While the ability to perform this coordinate transform in principle means the resulting beta weights are arbitrarily defined, they remain uniquely and informatively defined *given an understanding of the original model*. This fact should not be used to label model-based estimates as “arbitrary”, but instead emphasizes the importance of understanding the model used to derive conclusions about a dataset.

## Differences between conditions are preserved across linear transforms of the basis

At a high level, the IEM technique is a form of model-based dimensionality reduction. This approach estimates a transform from idiosyncratic measurement space (e.g. activation in voxels in V1; alpha power at EEG scalp electrodes) into a principled, manipulable, model-based “information” space. Perhaps most importantly, many studies using IEMs seek to compare channel response profiles, or basis-weighted ‘image’ reconstructions, across task conditions or timepoints in a trial. As described by Sprague et al (2018), these studies employ a *fixed encoding model*, such that activation patterns from different conditions are transformed into the same modeled information space, using a single common estimated encoding model (and often that encoding model is estimated using data from a completely different training task, e.g. Sprague et al., 2014, 2016, 2018b). In this case, the criticisms raised by Liu et al (2018) and Gardner & Liu (2019) do not apply: any arbitrary linear transforms would be applied equivalently to the results from each condition; and differences between conditions would be transformed from participant- and stimulus-specific measurement space into the same model-based ‘information’ space. Invertible transforms would serve only to adjust the axes of the modeled information space, providing a different ‘view’ of the same data. (Note that there may be cases where a transform renders differences between conditions invisible, but this would be exceedingly rare in cases where stimulus features span a feature space)

To make more concrete the point that differences between conditions can be preserved across linear transforms of the basis, we simulated an fMRI dataset for an experiment that contained two conditions, with one condition evoking a multiplicatively-larger response at the underlying neural level than the other (e.g., contrast, as in Liu et al., 2018; code [to be] available at: github.com/JohnSerences/iem_sim or github.com/tommysprague/iem_sim). Briefly, the response of each of 100 simulated voxels was computed as the sum of the responses of simulated neurons within each voxel, with each simulated neuron having a circular Gaussian tuning function across the feature space (with pseudo-randomly determined tuning bandwidth and amplitude, Figure 1A). The activity of the neurons within each voxel was computed in response to a set of 8 stimulus orientations across two experimental conditions, with multiplicative gain applied to the simulated neural responses in Condition 2 compared to Condition 1. One half of the data, balanced across stimulus type and experimental condition, were designated as a training set and the other half of the data were designated as a testing set. Using data in the training set, we next fit the voxel-wise forward encoding model comprised of 8 basis functions that span the feature space using either a standard set of raised cosine basis functions, tuned to specific feature values spanning the orientation space, or a set of raised cosine basis functions that were linearly transformed via an appropriately designed matrix into bimodal basis functions (termed the ‘xform’ matrix, Figure 1B; mirroring Gardner & Liu’s [2019] Fig. 2; the ‘P’ matrix in their notation). We then inverted both forward models, and used those inverted encoding models to reconstruct channel response profiles from the *same held-out test data*. Within each condition, channel response profiles mirrored the basis function used to estimate the corresponding model (Figure 2C, mirroring Gardner and Liu’s Figure 3). However, even though the shape of the channel response profiles is constrained predictably by the choice of the basis functions, differences between conditions are preserved: Condition 2 shows larger-amplitude channel response profiles regardless of the basis used. Importantly, because the transformation is linear and invertible, the bimodal channel response profiles from each condition can be losslessly converted back into unimodal channel response profiles via multiplication with the inverse of the original transformation matrix (Figure 1C, and note that this holds across a variety of gain modulations and with noise added at the level of simulated neurons, Figure 1D). Thus, one can apply arbitrary linear transforms to the basis set, and rather than rendering the data ‘arbitrary’, they remain interpretable *given knowledge of the encoding model*.

**Figure 1.**
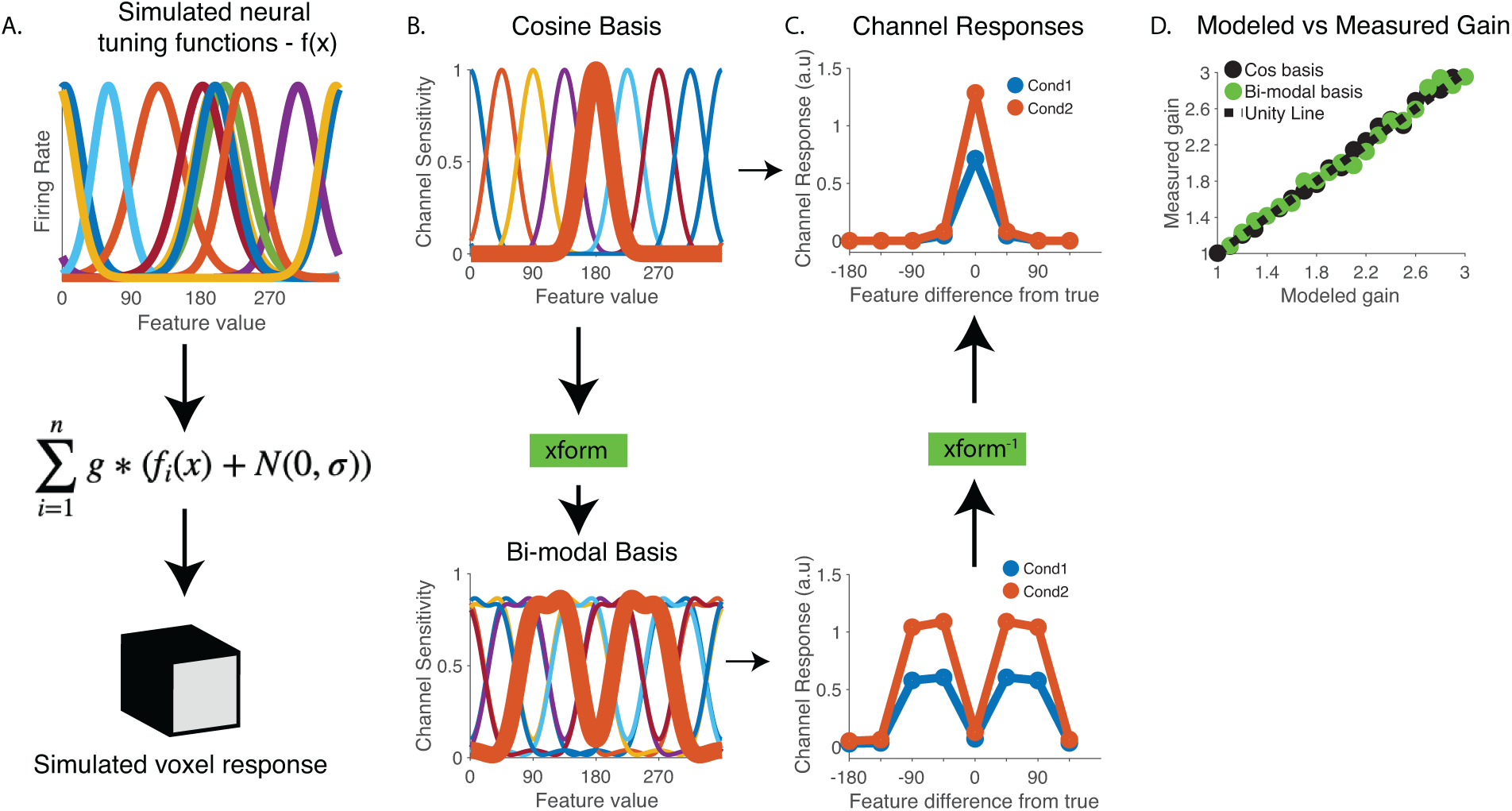
Differences between conditions can be preserved across invertible linear transforms. (A) We simulated voxel-level fMRI data where each voxel’s response was generated based on the sum of simulated responses across a population of simulated neurons with randomly centered tuning preferences (here, n=number of neurons, set to 100, although only 10 neural tuning functions are shown for clarity). Noise is added to the neural responses and then the gain factor (g) was applied to the data from each condition (Condition 1: g = 1, Condition 2: g = 1.8). For display purposes the noise (N) was set to 0 for panels A-C (following Fig. 3 in Gardner & Liu, 2019) and was set to 10 for panel D. (B) We analyzed data using two different formats of channel basis functions, mirroring those used by Gardner & Liu (2019). Importantly, the two bases are related by an invertible linear transform (xform). (C) Reconstructed channel response profiles differ in similar ways – Condition 2 has a higher ‘amplitude’ than Condition 1 – regardless of the basis set used, and the bi-modal channel response profiles are related by the inverse of the linear transform that was used to create the bimodal basis in the first place (xform^-1^). (D) Modeled gain compared to measured gain between Conditions 2 and 1, computed using both the raised cosine basis set and the transformed bi-modal version of the cosine basis set. Because there is not a straightforward way to quantify ‘amplitude’ for the channel response profiles computed from the bimodal basis, we instead implemented a model-free quantification scheme in which we computed the ratio of the area under each channel response profile (i.e. ratio of area under the curve in Condition 2 compared to Condition 1).

As shown in Figure 1, even though the shape of the channel response profiles is different due to the application of an invertible linear transform, the difference between conditions is preserved. This follows from the fact that, because the end result of the IEM procedure is a linear mapping from signal space into channel space, some differences in measured signals can be detected even across arbitrary basis transforms.

Thus, if the goal is to determine whether the amplitude of the channel response profiles increased, then the application of an invertible linear transform should not impact the general conclusions. Of course, this is true so long as one can accurately quantify or parameterize the resulting shape of the channel response profiles, which may be difficult if a random or oddly-shaped basis is used. Similarly, the process of aligning or re-centering channel response profiles on the correct feature can become vaguely defined if poorly motivated basis functions are used: typically, a unimodal channel is centered at the feature value to which it is tuned; but a bimodal or other oddly-shaped channel cannot be easily related to a particular feature value, further rendering data presentation and interpretation tricky in such cases. But again, we emphasize that, when channel response profiles are interpreted within the context of the model used to compute them, there is no sense in which the reported result is arbitrary.

## Comparison of IEM and Bayesian approaches to stimulus decoding

Gardner and Liu (2019) also make several other points. First, they highlight many positive aspects of the Bayesian decoding approach introduced by van Bergen et al (2015). We agree - van Bergen et al’s (van Bergen et al., 2015; van Bergen and Jehee, 2018) use of a forward model combined with a Bayesian readout rule is an innovative and promising technique, and thoughtfully analyzing data in different ways, especially when employing complex models, is always a good idea. In particular, the Bayesian decoding approach can provide complementary information about the uncertainty with which the activation pattern represents a feature value using an independently-estimated noise model, which is especially useful when trying to link trial-by-trial readouts of neural uncertainty with behavioral measures (van Bergen et al., 2015).

However, there are scenarios when directly comparing responses of modeled information channels can be informative: for example, Brouwer & Heeger (2011) compared responses at specific channels across contrast and stimulus conditions to evaluate the impact of cross-orientation suppression, and Ho et al (2012) and Scolari et al (2012) compared responses in channels tuned nearby the stimulus orientation across task (emphasize speed vs accuracy) and attention (target left vs target right) conditions. These types of analyses require estimating a full response profile across modeled channels, which is not easily accomplished with decoding analyses that generate a point estimate of the most likely stimulus feature (with or without a corresponding estimate of uncertainty). Moreover, when trying to disentangle responses associated with simultaneously presented stimuli, specifying an appropriate model in the Bayesian framework is not necessarily straightforward.

We also believe that each method, as well as other modeling approaches in this general domain, such as representational similarity analysis (RSA; Kriegeskorte et al., 2008; Kriegeskorte and Kievit, 2013), detailed voxel-wise encoding modeling using naturalistic stimuli (Kay et al., 2008; Naselaris et al., 2009; Nishimoto et al., 2011; Huth et al., 2012, 2016; Çukur et al., 2013a; Lescroart and Gallant, 2019), and simplified voxel RF modeling using focused stimulus sets (Dumoulin and Wandell, 2008; Serences et al., 2009; Saproo and Serences, 2010; Brouwer and Heeger, 2013; Wandell and Winawer, 2015; Mackey et al., 2017; Vo et al., 2017a), should be thoughtfully used to provide different insights into how information is encoded across a variety of task and stimulus conditions.

## Units of channel response profiles

Gardner and Liu (2019) also point out that the units of model-based reconstructions are arbitrary. This is a point that was noted in one of the original papers to use an IEM (Brouwer and Heeger, 2011). We agree that – reconstructed channel response levels are in arbitrary units, and we recommend researchers report them as such going forward. This, combined with the use of unit-normalized modeled channels (i.e., those used to predict channel responses when fitting the forward model), will render channel response estimates more comparable across studies. That said, it is essential to note that these units have no impact on the inferences that can be drawn when comparing channel response functions *between conditions* under a fixed encoding model. Thus, Gardner and Liu’s (2019) concerns about the arbitrary nature of this scale are not particularly germane to the interpretation of such results: one could scale all units by 42 without impacting the difference between conditions. Thus, so long as all model-based reconstructions that are compared head-to-head are on the same initial footing, then the comparisons are valid regardless of the conventions used to label the units of this analysis. However, when different models are trained for different conditions, it is less certain how to interpret differences in reconstructed channel response profiles across conditions: did the best-fit model, fit individually to each condition, change? Did the data used to reconstruct channel response profiles change? Did both change? By holding at least one aspect constant (the model, estimated with a neutral task or in a balanced fashion across conditions), it is possible to better ascertain how certain properties of neural response patterns change based on stimulus or task conditions as the units can be compared on equal footing (Sprague et al., 2018a).

## IEMs, and other analyses applied to voxel-based measurements, cannot be used to infer properties of single-unit tuning

Finally, Gardner and Liu (2019) and Liu et al (2018) imply that one of the goals of the IEM is to make inferences about single neuron response properties. Making inferences about the response properties of single-neurons is not possible using the IEM or any related model that operates at the scale of aggregate neural signals such as voxels, as different types of single-unit modulations can give rise to identical modulations at the level of a voxel (Sprague et al., 2018a). Thus, making such inferences is not the goal of the IEM or related measures, including the Bayesian decoding approach of van Bergen et al (2015). Instead, a fundamentally different approach that likely requires adopting a different measurement/analysis paradigm, such as parallel measurement of response properties measured across different scales (e.g., fMRI BOLD signal and single-unit electrophysiology; Keliris et al., 2019) would be needed to overcome the ill-posed many-single-neurons-to-voxel mapping problem.

## Defining terms

In the spirit of Gardner and Liu’s (2019) and Liu et al’s (2018) efforts to delineate the appropriate uses of IEMs, we want to more precisely define several terms related to the IEM technique to help clarify future reports. The IEM technique involves estimating an *encoding model* that best accounts for observed voxel activation responses given stimuli that are transformed into a modeled ‘channel space’ (and under the assumption of linearity such that the response of a given voxel is a linear combination of each of several modeled channels). Once an encoding model is estimated separately for each voxel, that encoding model can be *inverted* and used to *reconstruct* channel response profiles given new measured activation patterns across those same voxels. Those activation patterns are often measured in response to some kind of stimulus (either visual, or something attended, or held in working memory), and the resulting reconstructed channel response profiles typically contain representations of the stimulus/stimuli. To be clear, the result is not strictly a ‘stimulus reconstruction’, but a model-based reconstructed channel response profile. As an example, reconstructed channel response profiles for stimulus orientation are not literally an oriented grating. Instead, they describe the activation of modeled channels in response to a given stimulus, and this description is in a stimulus-referred space. Reconstructed channel response profiles can be used for several purposes, including *decoding* (recovering the most likely feature value(s) represented, and/or, with the use of an appropriate noise model, their uncertainty) and *quantification* (characterizing the shape of the channel response profile, including ‘width’, ‘amplitude’, etc., which should never be confused with the width or amplitude of single-neuron responses). Of course, all quantification of channel response profiles must be considered in concert with the encoding model used, but if a fixed encoding model is used for reconstructing channel response profiles across several experimental conditions, their properties can be compared in the context of the model.

## Conclusions

In this reply to Gardner and Liu (2019) commentary, we hope to have clarified some mischaracterizations of how the IEM approach is carried out (see also: Sprague et al., 2018a). To be clear, we are not arguing that the IEM or related approaches are not without serious limitations - the model specification is key, as is understanding what inferences can and cannot be supported by the results (Sprague et al., 2015, 2018a). As Gardner and Liu (2019) point out, these limitations are especially important to recognize when modelling signals in feature spaces that are not well-understood, such as those for complex shapes or for higher-order cognitive or social functions. In these situations, an IEM may still be able to quantify differences between conditions and could thus be used to make inferences about changes in the information content of population-level response patterns. However, drawing links between the shape of IEM-derived channel response profiles and the properties of population-level neural representations is not appropriate. Instead, we agree with the suggestions of Gardner & Liu (2019) that careful comparison of forward models that are not related by an invertible linear transform is better suited for this purpose (e.g., Brouwer and Heeger, 2009; Nishimoto et al., 2011; Lescroart and Gallant, 2019). That said, IEM-based channel response profiles are not arbitrary when the model choice is based on principled assumptions about neural population representations and, more importantly, channel response profiles are uniquely determined given knowledge of the modeled basis, whatever that basis may be.

We believe the IEM method is most useful when comparing reconstructed channel response profiles across manipulations of stimulus properties (e.g., contrast) or task conditions (e.g., attention), or combinations thereof (Sprague et al., 2018b) using a fixed encoding model across relevant comparisons (Sprague et al., 2018a). When used this way, the criticisms raised by Gardner & Liu (2019) have no substantial bearing on the efficacy of the IEM technique for comparing the impact of experimental manipulations on information represented within aggregate measurements of neural activity patterns. In other words, in the same way the answer (42) is only meaningful in the context of the Question, results derived from a model are only meaningful in the context of the model used.

## Acknowledgements

Funding provided by NEI-EY025872 to JTS. We thank Kirsten Adam, Edward Awh, Clayton Curtis, Barry Giesbrecht, Margaret Henderson, and Bradley Postle for helpful discussions and comments on the manuscript.

## Footnotes

1 We note that Gardner & Liu (2019) use “channel response function” in their commentary, and others have used “channel tuning function”; we elect to refer to results from the IEM technique as channel response profiles, to further distance these results from single-neuron tuning functions.

